# A Novel PD-L1 Splice Isoform Modulates β Cell Communication in Response to Interferon Signaling

**DOI:** 10.64898/2026.09.25.754099

**Authors:** Chaitra Rao, Fei Huang, Saptarshi Roy, Matthew C. Austin, Matthew B. Johnson, Kerim B. Kaylan, Irene Amalraj, Andre G. De Oliveira, Decio L. Eizirik, Carmella Evans-Molina, Amelia K. Linnemann, Jon D. Piganelli, Richard Oram, Raghavendra G. Mirmira, Emily K. Sims

## Abstract

β cell expression of the immune checkpoint ligand PD-L1 (encoded by *CD274)* limits autoimmune β cell destruction in type 1 diabetes (T1D). β cell display PD-L1 not only at the cell surface but also secreted on extracellular vesicles (EVs), which bind PD-1 and restrain CD8^+^ T cell activation. β cell IFN signaling is an early driver of T1D pathogenesis, yet the mechanisms linking IFN signaling to the fate of PD-L1 protein remain undefined. Here, we identify an IFN- and coxsackievirus-inducible alternatively spliced isoform of PD-L1, PD-L1^Δ3^, lacking cassette exon 3 and generated in human β cells and islets in response to IFN signaling or viral infection. PD-L1^Δ3^ transcripts are elevated in islets from donors with single autoantibody positivity (AAB^+^) and T1D. Unlike full-length PD-L1, which localizes to the plasma membrane, PD-L1^Δ3^ is retained intracellularly and loses the capacity to bind PD-1. Functionally, PD-L1^Δ3^ fails to suppress CD8^+^ and CD4^+^ T cell proliferation, activation, and cytotoxic cytokine release, and is not efficiently sorted into EVs. Convergently, a germline *CD274* splice-site variant (c.682+1G>A) found in siblings with neonatal T1D yields a protein with the same intracellular retention, reduced β cell PD-1 binding, and reduced β cell and circulating EV PD-L1. Together these findings define PD-L1^Δ3^ as an IFN-induced splice variant that diverts PD-L1 away from its immunoregulatory, EV-competent form, revealing a post-transcriptional axis that shapes β cell immune communication.

**Article highlights:**

- IFNs and coxsackievirus B induce a novel exon 3-skipped PD-L1 isoform (PD-L1^Δ3^) in human β cell and islets.
- PD-L1^Δ3^ transcripts are elevated in islets from human donors with AAB+ and T1D.
- PD-L1^Δ3^ is retained intracellularly, does not bind PD-1 and fails to suppress T cell responses.
- PD-L1^Δ3^ is excluded from EVs, diverting PD-L1 from its immunoregulatory, EV-competent form.
- A germline *CD274* splice variant (c.682+1G>A) in siblings with neonatal T1D reproduces these defects and lowers circulating EV PD-L1.

## Introduction

Type 1 diabetes (T1D) is among the most common chronic autoimmune diseases of childhood and represents a major and rapidly growing global health burden^1^. Current estimates indicate that over 9 million individuals live with T1D, including approximately 1.8 million children and adolescents, and incidence continues to rise worldwide^2^. This figure likely underestimates the true burden, as adult-onset autoimmune diabetes is frequently misclassified as type 2 diabetes^3,4^. While immunotherapies and improved metabolic management can delay β cell decline, clinical responses remain highly variable^5–7^. This heterogeneity reflects a fundamental gap in our understanding of factors driving disease progression, and more specifically, an incomplete understanding of how β cell-intrinsic signaling pathways, ranging from stress-adaptive to maladaptive programs, regulate immune engagement and β cell survival^8–10^.

Intercellular communication at the β cell–immune cell interface is a likely early contributor to T1D progression^11^. A key immune regulatory mechanism involves β cell expression of the checkpoint protein programmed death ligand 1 (PD-L1), which engages with its receptor programmed death-1 (PD-1) on immune cells to suppress activity and limit autoimmune destruction^12^. Donors with long-standing T1D exhibit increased PD-L1 in insulin-positive β cells, while PD-L1 is absent in insulin-deficient islets^13^, suggesting that upregulation is associated with insulin positive β cell survival. Recent evidence from our group demonstrated that β cells do not only express PD-L1 locally but also export PD-L1 on the surface of extracellular vesicles (EVs)^14^. These PD-L1+ β cell EVs have been shown to bind PD-1 and suppress CD8+ T cell proliferation and cytotoxicity, extending the immunoregulatory reach of the β cell beyond direct cell-cell contact.

Intracellularly, β cell IFN signaling not only alters transcript abundance but also impacts RNA processing, including alternative splicing, an emerging pathway linking cellular stress to immune dysfunction^15–17^. Alternative splicing enables a single gene to generate multiple transcript isoforms, expanding proteomic diversity and altering protein function in response to inflammatory cues^18^. Cytokine exposure remodels β cell splicing machinery and splicing events in human islets have been shown to contribute to β cell dysfunction and neo-epitope generation in T1D^19–23^. Alternative splicing events in whole blood are also enriched in individuals with new-onset T1D and distinguish them from unaffected controls^21^. In cancer, *CD274,* the gene encoding PD-L1, undergoes alternative splicing to generate multiple isoforms, including a variant lacking exon 3, which have been reported to play distinct roles in immune surveillance and disease progression^24–29^. Whether IFN signaling drives comparable splicing of *CD274* in β cells, and whether the resulting isoforms differ in trafficking or fate of PD-L1 remains undefined. Specifically, the mechanisms by which IFN-induced signaling directs the partitioning of PD-L1between the cell surface and incorporation as EV cargo are unknown.

Here, we address these gaps by investigating the molecular link between early β cell IFN signaling and the β cell’s subsequent capacity to maintain immune tolerance. We hypothesize that IFN signaling alters not only the amount of PD-L1 produced by β cells, but also which isoform is made and whether it reaches the cell surface and EV compartment. By defining the mechanisms governing EV cargo selection and identifying how these responses reflect inter-individual heterogeneity, we uncover a novel post-transcriptional axis that shapes β cell–immune communication and that may provide a foundation for developing minimally invasive biomarkers to track β cell IFN signaling and alternative splicing.

## Results

### 1. Identification of a novel PD-L1 splice isoform (PD-L1^Δ3^) in human β cells and islets

To investigate the mechanisms promoting PD-L1 expression in the context of T1D, we first modeled the early inflammatory islet microenvironment using IFNs. IFNs are implicated in viral-mediated β cell stress and are known to elicit upregulation of PD-L1^30–33^. Treatment of primary human islets with IFN-□ resulted in a robust upregulation of PD-L1 expression. Notably, Western blot analysis revealed a dual-banding pattern in IFN-treated human islets. In addition to the expected full-length PD-L1 band (42-45 kDa), we detected a distinct, lower molecular weight (MW) band (∼37-41 kDa). The size of this smaller protein is consistent with the predicted MW of an alternatively spliced PD-L1 variant. Although induction of both PD-L1 bands was pronounced in whole-islet lysates, the lower-MW band was notably absent in EV fractions, suggesting a selective cellular retention of this isoform or a bias toward full-length PD-L1 for EV cargo loading (Figure 1A).

**Figure 1.**
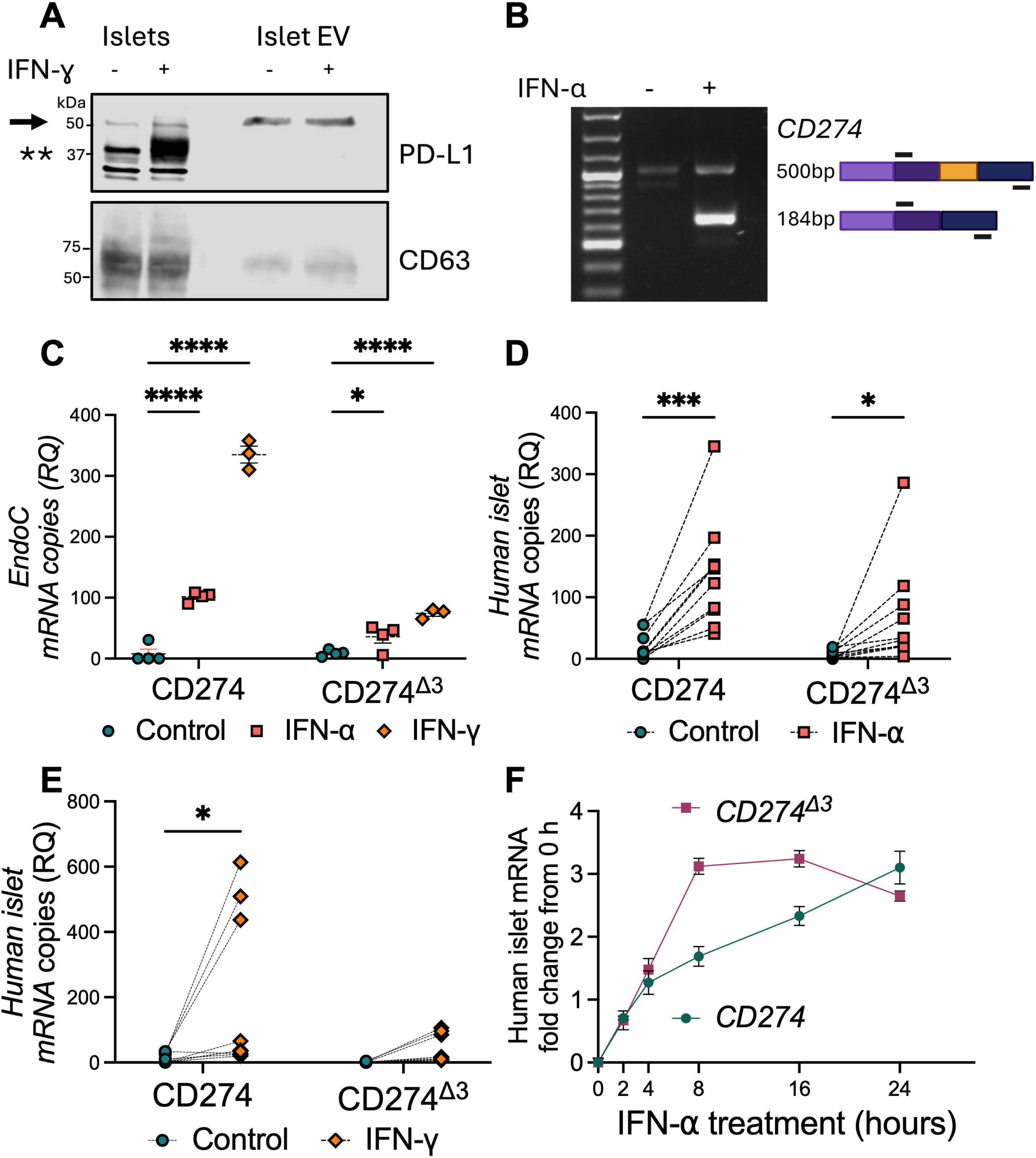
IFN signaling induces a novel exon 3-skipped PD-L1 isoform (PD-L1^Δ3^) in human β Cells and islets. (A) Immunoblot of PD-L1 and CD63 in whole islet lysates and islet derived EVs from primary human islets treated with 100 ng/µl IFN-□ for 24 h. Arrow, full-length PD-L1; double asterisk, lower-molecular weight PD-L1 (B) RT-PCR using primers spanning exon 2-4 in EndoC-βH1 cells +/- 2000 U/ml IFN-α, with schematic of the corresponding transcripts. (a-b) Representative of 3 experiments. (C-E) Isoform-specific qPCR for full-length CD274 and CD274^Δ3^ in (C) EndoC-βH1; N=3, human islets treated with either (D) IFN-α or (E) IFN-□ for 24 h, N=10. (F) Time course of CD274 and CD274^Δ3^expression in human islets +/- IFN-α from 0-24 h, N=3. RQ; relative quantity. Data represented as mean SEM. Statistical comparisons were performed by [paired 2-tailed t test / 1-way ANOVA with Tukey’s multiple-comparisons test]. * P <0.05, *** P <0.0005, **** P <0.0001

To characterize this variant at the transcript level, we performed RT-PCR using primers flanking the region between exon 2 and exon 4. This analysis confirmed the presence of a shortened transcript (184 bp) in addition to the full-length amplicon (∼500 bp), with the shortened product markedly enriched following IFN-α treatment (Figure 1B). Subsequent isoform-specific qPCR in both the EndoC βH1 human β cell line and primary human islets (Table 1) demonstrated that IFN-α or IFN-□ significantly upregulate both full-length *CD274* and the truncated isoform, hereafter PD-L1^Δ3^ (Figure 1C-E). A time-course analysis following IFN-α exposure revealed that PD-L1^Δ3^ induction is rapid, rising within the first hours of treatment with peak levels preceding the slower accumulation of full-length *CD274* (Figure 1F). These results establish PD-L1^Δ3^ as an interferon-inducible isoform whose kinetics are distinct from those of the full-length transcript.

**Table 1:** Non-diabetic donor islet characteristics.

| RRID | Age | BMI | Hba1c | Sex | Islet source |
| --- | --- | --- | --- | --- | --- |
| SAMN37350251 | 43 | 29.9 | - | Female | IIDP |
| SAMN36704819 | 52 | 29.9 | - | Male | IIDP |
| SAMN43813303 | 35 | 28.9 | 5.4 | Male | ADI core |
| SAMN44368156 | 56 | 31.3 | 5.6 | Male | ADI core |
| SAMN44368157 | 64 | 31.7 | 5.6 | Female | ADI core |
| SAMN44624610 | 54 | 27.3 | 5.6 | Female | ADI core |
| SAMN45037497 | 59 | 26.6 | 4.5 | Male | ADI core |
| SAMN45114368 | 37 | 28.7 | 5.4 | Male | ADI core |
| SAMN47415044 | 38 | 32.5 | 5.8 | Male | ADI core |
| SAMN47939314 | 34 | 24.2 | 5.1 | Male | ADI core |
| SAMN48086628 | 34 | 31.8 | 5.4 | Male | ADI core |
| SAMN48134749 | 64 | 27.3 | 5.7 | Male | ADI core |
| SAMN51754307 | 41 | 29.5 | 5.8 | Male | ADI core |
| SAMN52371583 | 18 | 22.7 | 5.1 | Male | ADI core |
| SAMN53736986 | 66 | 22 | 6.4 | Male | ADI core |
| SAMN47834917 | 39 | 22.9 | 5.7 | Male | IIDP |
| AMEW466 | 36 | 35 | - | Female | UNOS |
| SAMN47541014 | 45 | 31.9 | 5.6 | Male | IIDP |
| AMEC275 | 39 | 39 | - | Female | UNOS |
| AMA3262 | 44 | 23.7 | - | Male | UNOS |
| AMBC032 | 60 | 45 | 5 | Female | UNOS |

### 2. Coxsackievirus infection induces PD-L1Δ3 in human islets

We next asked whether this splicing event occurs under pathophysiologically relevant stress that would be expected to induce IFN signaling. Consistent with our cytokine models, coxsackievirus B (CVB) infection, a viral trigger strongly associated with T1D, also induced PD-L1^Δ3^ in human islets (Figure 2A). BaseScope for full-length PD-L1 and PD-L1^Δ3^ in insulin-positive cells showed that both isoforms increased following 24 hour CVB exposure, with PD-L1^Δ3^ integrated intensity significantly elevated in CVB-infected islets relative to controls, exhibiting a more pronounced increase than that of full-length PD-L1 (Figure 2A-B). Together with the IFN data, these results identify PD-L1^Δ3^ as a distinct splice variant that is transcribed alongside full-length PD-L1 in response to T1D-relevant triggers.

**Figure 2.**
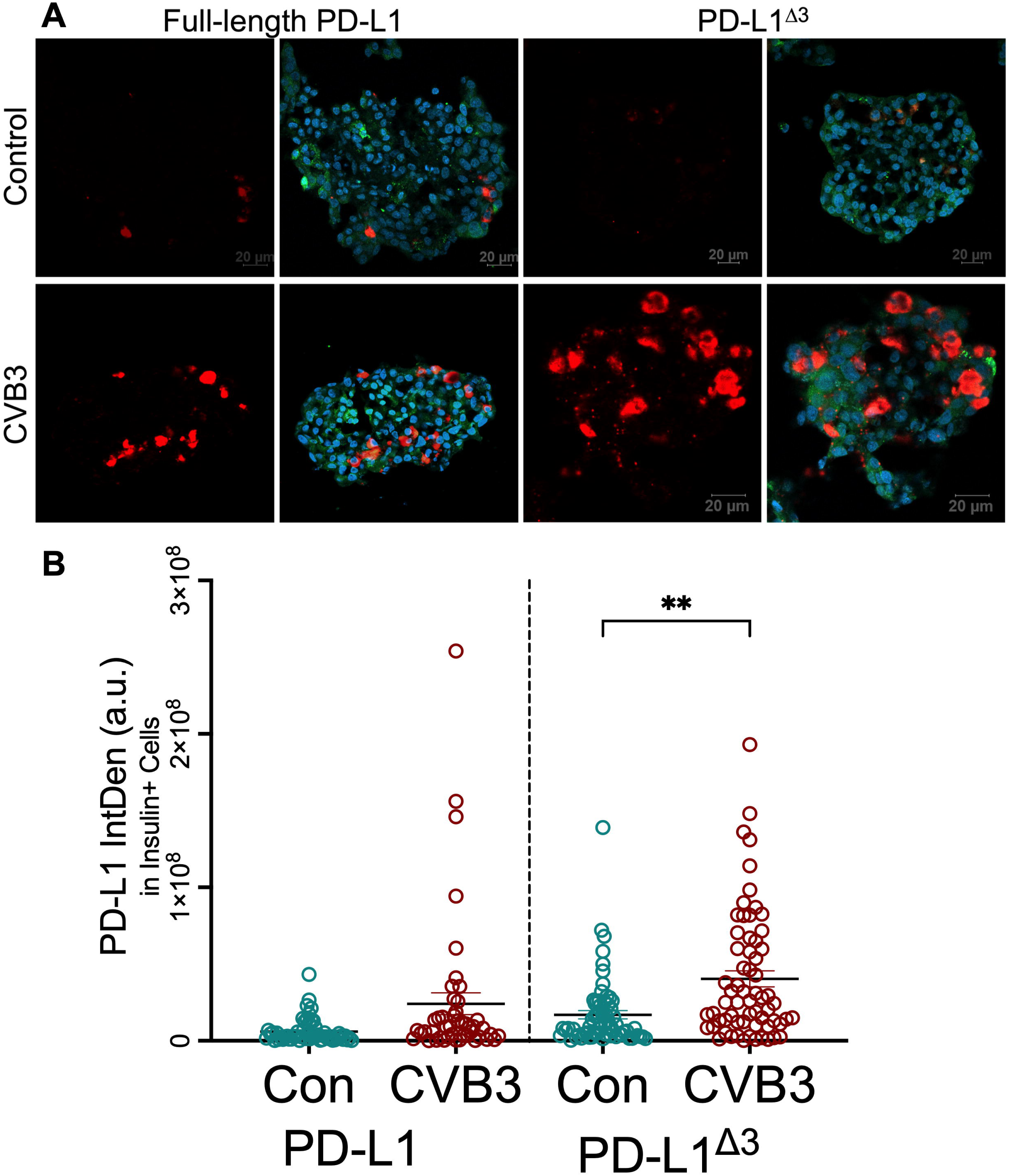
Coxsackievirus B infection induces PD-L1Δ3 in human islets. (A) Representative confocal images of BaseScope in situ hybridization of full-length PD-L1 and PD-L1Δ3 (red) with insulin (green) and DAPI (blue) in primary human islets 24 hr after mock (control) or coxsackievirus B (CVB3, MOI = 50) infection. Scale bars 20 µm. (B) Quantification of PD-L1 and PD-L1^Δ3^ integrated density within insulin positive cells. Each symbol represents islets from 6 independent donors per condition. Data represented as mean SEM. ** P <0.005.

### 3. Human islet PD-L1Δ3 is elevated in donors with T1D and AAB^+^

To determine whether PD-L1^Δ3^ is relevant to human disease, we quantified isoform-specific transcripts in insulin positive β cells in situ using BaseScope in pancreatic sections from non-diabetic (ND), single autoantibody-positive (AAB^+^), and donors with T1D (N=3/group) (Table 2). Full-length PD-L1 transcript abundance was low in ND islets and varied widely between donors. Transcript abundance progressively increased across AAB^+^ and T1D groups, consistent with prior reports of full-length PD-L1 upregulation in residual insulin positive β cells in T1D^13,34^ (Figure 3A-B). PD-L1^Δ3^ transcripts exhibited a similar disease-associated pattern, remaining near background in ND islets with elevated expression in AAB^+^ donor islets and residual insulin+ islets in donors with T1D. Even within disease groups, relative abundance of PD-L1^Δ3^ vs. full-length transcripts varied from donor to donor, with 1/3 donors within both the single AAB^+^ and the T1D groups exhibiting a higher fraction of PD-L1^Δ3^ vs. full-length PD-L1. (Figure 3C). Interestingly, CD3 immunostaining of the section from the donor in the T1D group with the high fraction of PD-L1^Δ3^ transcript numbers frequently identified islets with T cell infiltration (Figure 3D). Overall, these data confirm that β cell PD-L1^Δ3^ splicing is associated with human autoimmunity and T1D, appearing early after establishment of islet autoimmunity.

**Figure 3.**
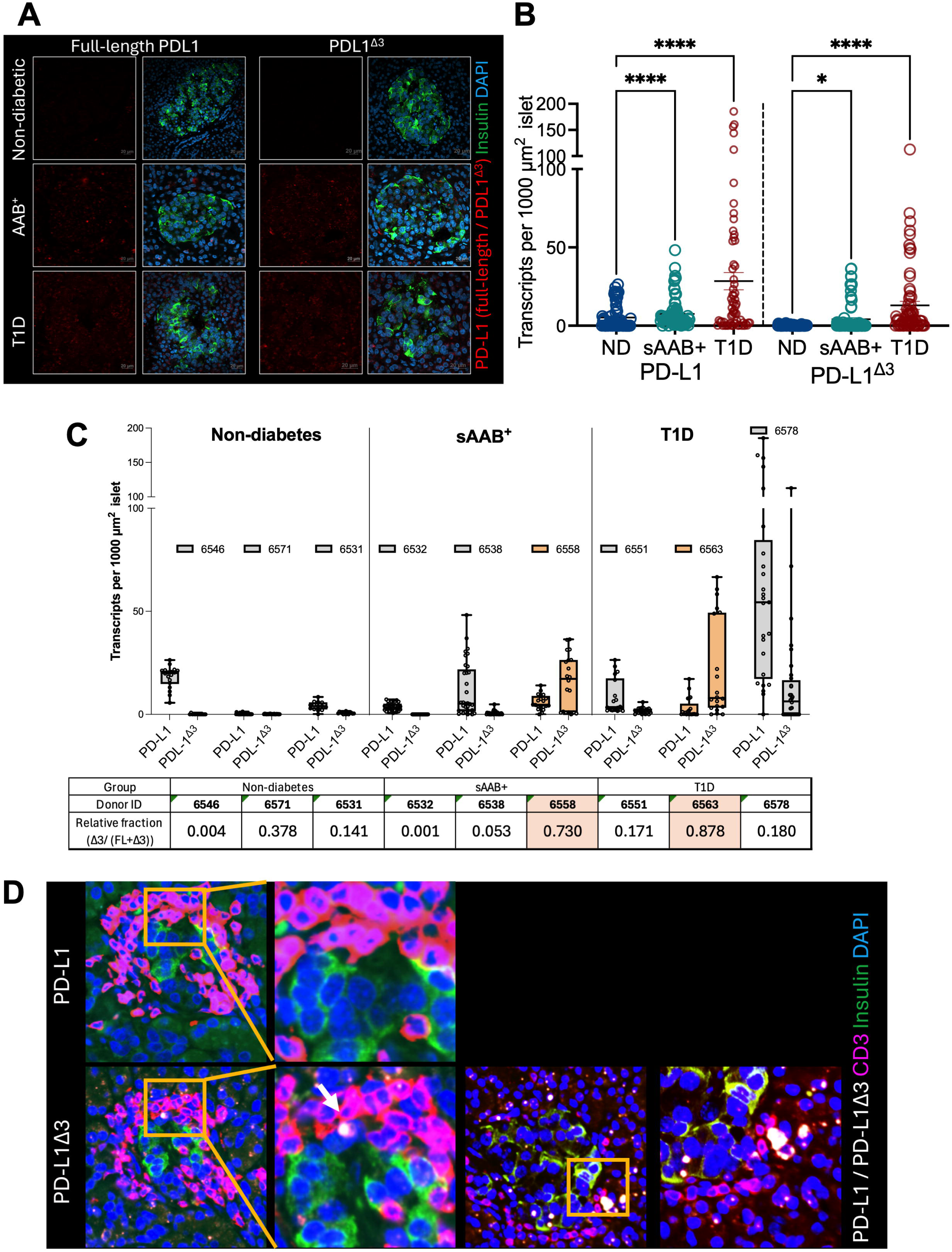
PD-L1^Δ3^ is elevated in human T1D and AAB+ islets. (A) Representative BaseScope in situ hybridization for full-length PD-L1 and PD-L1^Δ3^ (red) with insulin (green) and DAPI (blue) in pancreatic sections from non-diabetic (ND), single autoantibody positive (AAB^+^) and donors with T1D. A median of 20 islets was quantified per donor (range 13-44 islets per donor, N=3 donors per group). Scale bars: 20 µm. (B-C) Transcripts per 1000 µm^2^ islet area for full-length PD-L1 and PD-L1^Δ3^ (B) in all the donors per group (C) quantified by individual donor with relative PD-L1^Δ3^ fraction calculated per donor as [mean PD-L1^Δ3^ density÷(mean PD-L1^Δ3^ density + mean full-length PD-L1 density)]. (D) BaseScope in situ hybridization for full-length PD-L1 and PD-L1^Δ3^ (white) with insulin (green). CD3 (pink) and DAPI (blue) in donor with high fraction of PD-L1^Δ3^. * P <0.05, **** P <0.0001

**Table 2:** Human pancreatic tissue donor characteristics.

| **nPOD caseID** | **RR_id** | **donor_type** | **sex** | **race_ethnicity** | **age_years** | **GADA** | **IA_2A** | **mIAA** | **ZnT8A** |
| --- | --- | --- | --- | --- | --- | --- | --- | --- | --- |
| 6531 | SAMN18053213 | No Diabetes | Female | Hispanic/Latino | 19.25 | - | - | - | - |
| 6532 | SAMN18053214 | Autoantibody + | Male | Hispanic/Latino | 20.04 | + | - | - | - |
| 6538 | SAMN25652249 | Autoantibody + | Male | Caucasian | 19.14 | + | - | - | - |
| 6546 | SAMN25652257 | No Diabetes | Male | Asian | 22.29 | - | - | - | - |
| 6551 | SAMN25652262 | Type 1 Diabetes | Male | Caucasian | 20.7 | + | + | + | + |
| 6558 | SAMN30386847 | Autoantibody + | Female | African American | 21.69 | + | - | - | - |
| 6563 | SAMN30386851 | Type 1 Diabetes | Female | Caucasian | 14.56 | - | + | - | - |
| 6571 | SAMN33284289 | No Diabetes | Male | African American | 28.1 | - | - | - | - |
| 6578 | SAMN33284295 | Type 1 Diabetes | Female | Caucasian | 11.95 | - | + | - | + |

### 4. Intracellular localization of PD-L1^Δ3^ in human β cells and loss of PD-1 binding capacity

To investigate the functional consequences of the PD-L1 isoform shift, we generated human β cells overexpressing tagged versions of each isoform, taking advantage of their minimal endogenous PD-L1 expression. Flow cytometry for surface PD-L1 confirmed robust cell-surface presentation of full-length PD-L1, whereas surface staining for PD-L1^Δ3^ was comparable to that of the empty-vector, indicating minimal plasma-membrane display despite abundant intracellular expression (Figure 4A-B). Subcellular fractionation confirmed that full-length PD-L1 partitioned predominantly to the membrane fraction, while FLAG-tagged PD-L1^Δ3^ was enriched in the cytoplasmic fraction, with both isoforms largely excluded from the nucleus (Figure 4C-D).

**Figure 4.**
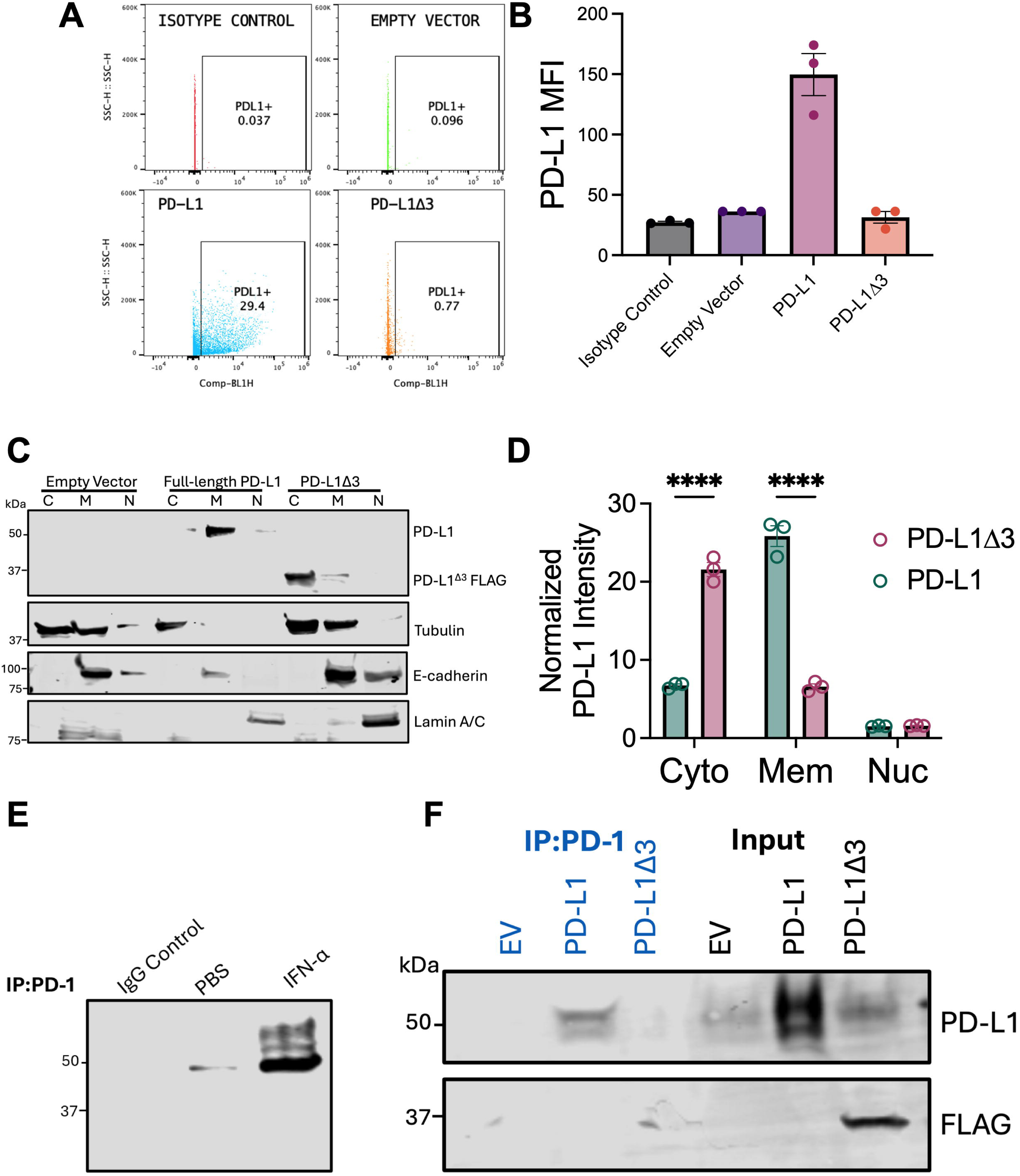
PD-L1^Δ3^ is retained intracellularly and does not bind PD-1. (A-B) Representative flow cytometry plots and quantification of surface PD-L1 mean fluorescence intensity (MFI) in 3 independent experiments; each symbol represents 1 experiment. N=3 (C-D) Immunoblot of cytoplasmic (C), membrane (M), and nuclear (N) fractions from human βlox-5 cells expressing full-length PD-L1 or FLAG-tagged PD-L1^Δ3^, with tubulin, E-cadherin and lamin A/C as fraction-purity controls, and quantification of normalized PD-L1 intensity per fraction. (E) PD-1 pulldown from βlox-5 cell lysates treated +/- IFN-α, followed by immunoblot for PD-L1. Bead-only control included. (F) PD-1 pulldown from lysates of cells expressing empty vector (EV), full-length or PD-L1^Δ3^, with corresponding input, immunoblotted for PD-L1 and FLAG. N=3 independent experiments. Data represented as mean SEM. **** P <0.0001

Because exon 3 lies within the immunoglobulin-variable-like (IgV-like) PD-1-binding interface, we also asked whether the retained, intracellular isoform could still engage PD-1. Using a PD-1 pulldown assay in β cell lysates, IFN-α treatment increased the recovery of β cell PD-L1 on PD-1 beads, confirming inducible PD-1 engagement of full-length PD-L1 (Figure 4E). In overexpression lysates, only full-length PD-L1 and not PD-L1^Δ3^ was recovered by PD-1 pulldown, demonstrating that PD-L1^Δ3^ loses intrinsic PD-1 binding capacity (Figure 4F). Thus, PD-L1^Δ3^ is both trafficking-defective and binding-defective: it is largely retained in the cytoplasm rather than displayed at the cell surface, and even overexpressed protein cannot engage PD-1 effectively, positioning it poorly to restrain immune cells.

### 5. PD-L1^Δ3^ fails to suppress T-cell proliferation and cytotoxicity

Given that PD-L1^Δ3^ is intracellularly retained and cannot bind PD-1, we tested the immunoregulatory consequences of a PD-L1^Δ3^ isoform shift by co-culturing stimulated human T cells with β cells overexpressing full-length PD-L1 or PD-L1^Δ3^ or empty vector and tracking responses over 72 h. Full-length PD-L1 markedly suppressed both CD8^+^ and CD4^+^ T cell proliferation, as measured by CellTrace Violet dilution, restoring proliferation toward unstimulated baseline levels. In contrast, PD-L1^Δ^ ^3^-expressing β cells exhibited significantly reduced capacity to suppress proliferation (Figure 5A-B). Time-course analysis showed increased CD8^+^ and CD4^+^ T-cell proliferation over 72 h in the stimulated, empty-vector, and PD-L1^Δ3^ groups, while proliferation remained low with full-length PD-L1 and in unstimulated cells (Figure 5C-D). Overall viability across conditions was similar (>85% live), indicating that these differences reflect suppression rather than cytotoxicity toward the T cells (Figure 5E).

**Figure 5.**
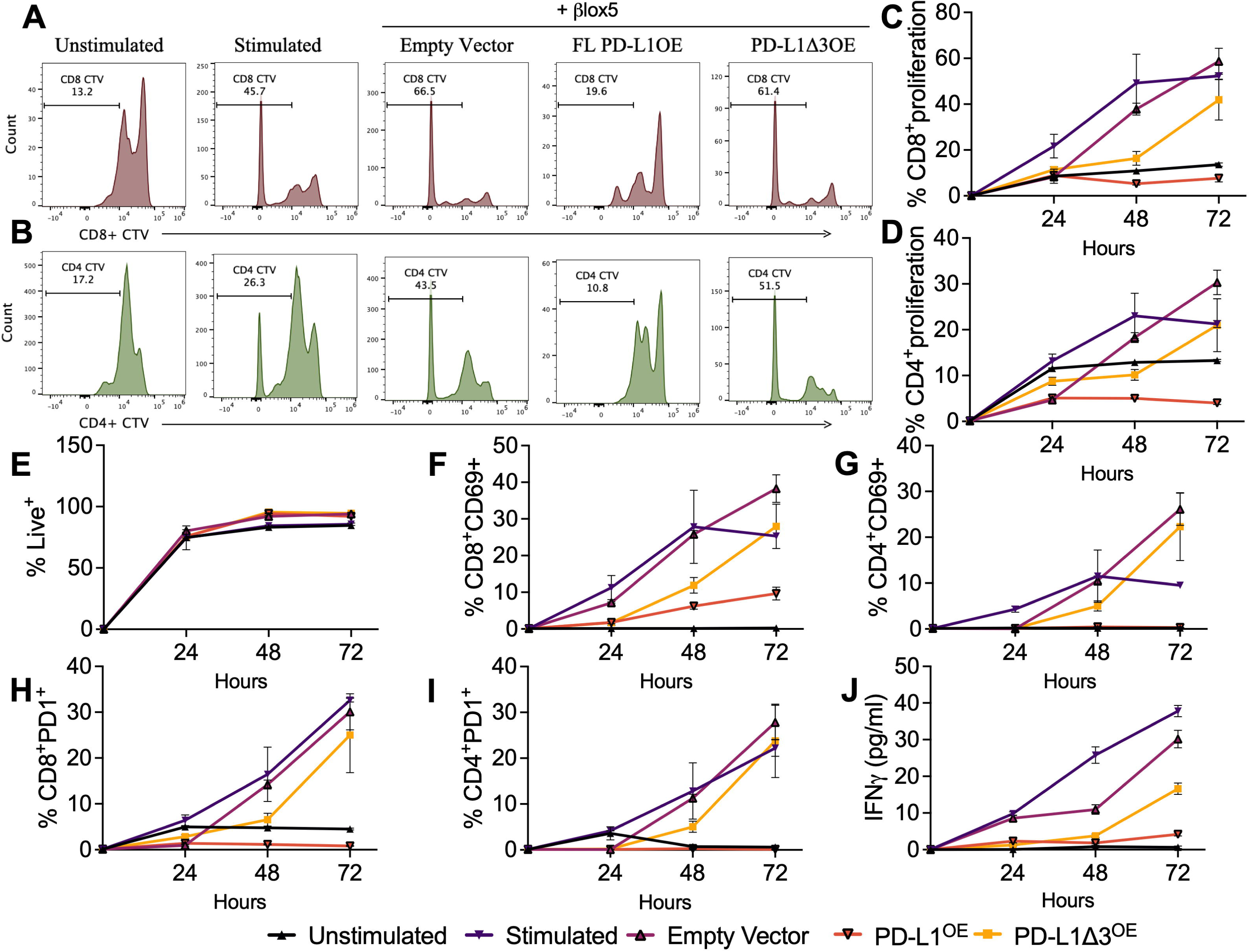
PD-L1^Δ3^ fails to suppress T-cell proliferation, activation and IFN-□ secretion. Human PBMCs were labeled with CellTrace Violet (CTV), stimulated, and co-cultured for up to 72 h with human βlox5 cells expressing empty vector, full-length PD-L1 or PD-L1^Δ3^. (A-B) Representative CTV dilution for (A) CD8^+^ and (B) CD4^+^ T cells at 72 h. (C-D) Quantification of (C) CD8^+^ and (D) CD4^+^ T cell proliferation over time. (E) Percentage of live cells. (F-G) Percentage of (F) CD8^+^CD69^+^ and (G) CD4^+^CD69^+^ T cells. (H-I) Percentage of (H) CD8^+^PD1^+^ and (I) CD4^+^PD1^+^ T cells. (J) IFN-□ concentration in co-culture supernatants measured by ELISA. N=3 independent experiments using PBMCs from 2 non-diabetic donors. Data represented as mean SEM.

Consistent with impacts on proliferation, full-length PD-L1 blunted T cell activation, reducing the frequencies of CD8^+^CD69^+^ and CD4^+^CD69^+^ cells. In contrast, PD-L1^Δ3^ conferred only partial inhibition of CD69 induction, incompletely restraining T cell activation relative to full-length PD-L1 (Figure 5F-G). Surface PD-1 on CD8^+^ and CD4^+^ T cells was reduced in the presence of full-length PD-L1, consistent with successful ligand engagement and receptor downmodulation; PD-L1^Δ3^ produced no such reduction, in line with its reduced ability to bind PD-1 (Figure 5H-I). Finally, full-length PD-L1 suppressed secretion of the pro-inflammatory cytokine IFN-γ, while PD-L1^Δ3^ cocultures showed only partial suppression of IFN-γ output (Figure 5J). Together, these data establish that compared to full-length PD-L1, PD-L1^Δ3^ exhibits impaired functional capacity to restrain T cell responses, consistent with its intracellular retention and reduced PD-1 binding.

### 6. PD-L1^Δ3^ does not sort to β cell EVs

Because β cells present immunomodulatory full-length PD-L1 on both plasma and EV membranes, we also tested whether PD-L1^Δ3^ is competent for EV loading using human β cells overexpressing full-length PD-L1 or PD-L1^Δ3^. To measure total EV PD-L1, we used a PD-L1 antibody directed against the cytoplasmic tail near the C-terminus, a region outside exon 3 and therefore retained in both isoforms. EVs from cells expressing full-length PD-L1 were strongly positive for total PD-L1 staining. In contrast, PD-L1^Δ3^ cell EVs carried substantially less total PD-L1 despite comparable tetraspanin content, showing an ∼5-fold lower total PD-L1 signal on PD-L1^Δ3^ EVs (Figure 6A-B). To quantify isoform partitioning independent of a single tetraspanin, we captured EVs on CD63, CD81, or CD9 and measured associated total PD-L1 across all three capture antibodies. Again, PD-L1^Δ3^ isoform exhibited approximately one-quarter (less that <20%) of the total EV-associated PD-L1 compared to the cells overexpressing full-length PD-L1 (Figure 6C). Thus, exon 3 skipping selectively excludes PD-L1 from the EV export pathway, depleting the pool of PD-L1+ immunoregulatory EVs.

**Figure 6.**
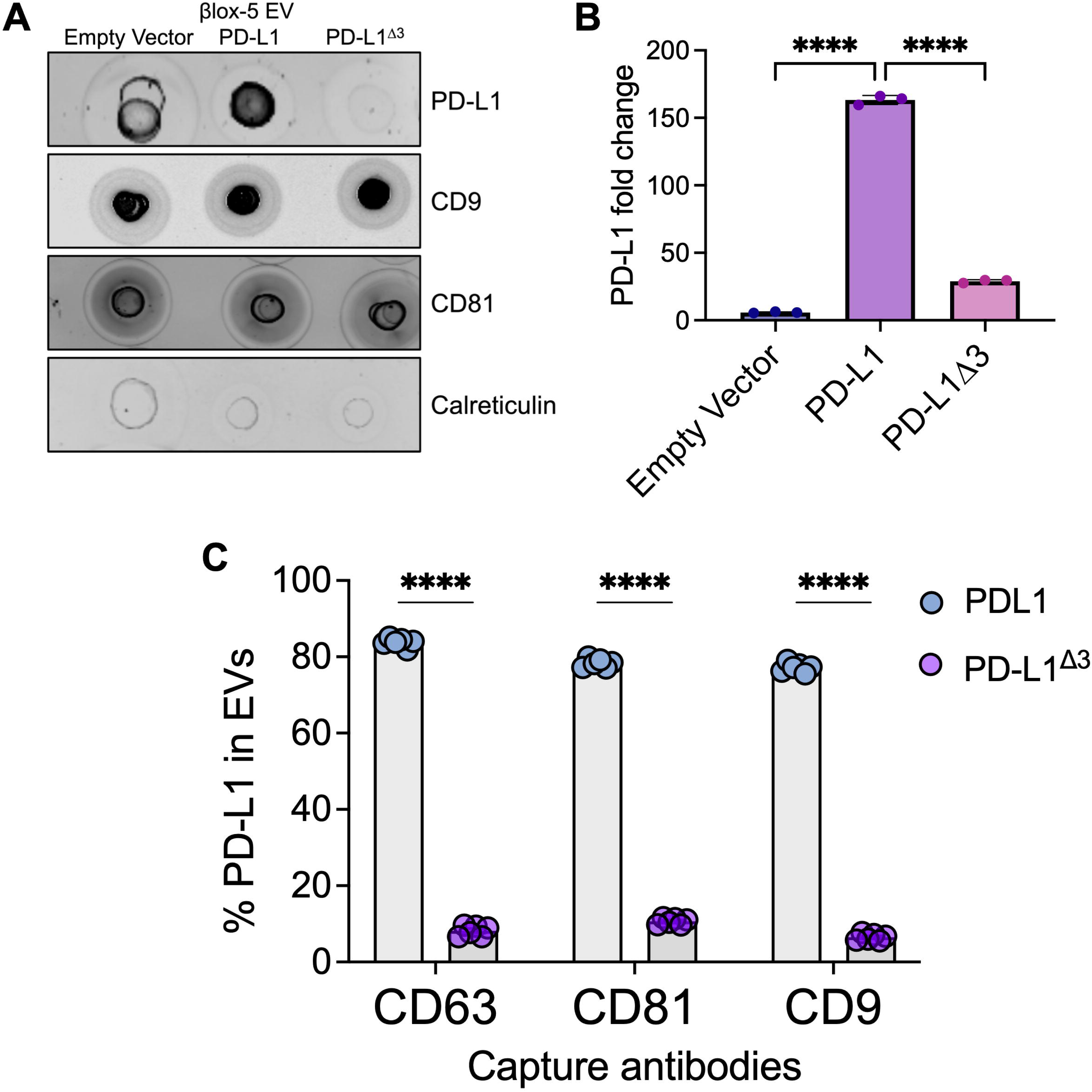
PD-L1^Δ3^ is not sorted into β cell EVs. (A) Dot blot for PD-L1, CD9, CD81 and calreticulin in EVs isolated from human βlox5 cells expressing empty vector, full-length PD-L1 or PD-L1^Δ3^. (B) Quantification of EV PD-L1 signal expressed as fold change relative to empty vector. (C) Percentage of EV-associated PD-L1 βlox5 cells expressing full-length PD-L1 or PD-L1^Δ3^ following EV capture on CD63, CD81 or CD9 tetraspanin antibodies. N=3 independent experiments. Data represented as mean±SEM. Statistical comparisons were performed by (B) [1-way ANOVA with Tukey’s multiple-comparisons test and (C) 2-way ANOVA with Šidák’s multiple-comparisons test (C)]. ****P < 0.0001.

### 7. A germline CD274 splice-site variant reproduces PD-L1^Δ3^ features and reduces circulating EV PD-L1

To test whether disrupted PD-L1 splicing compromises β cell: immune cell signaling in human genetics, we studied a family carrying a germline splice-donor-site variant in *CD274* (c.682+1G>A)^35^ resulting in neonatal autoimmune diabetes in two homozygous siblings (Figure 7A). The c.682+1G>A variant disrupts a canonical 5′ splice donor site predicted to alter constitutive splicing of *CD274*, providing a naturally occurring, genetically encoded example of aberrant PD-L1 splicing in humans. The putative PD-L1 protein was predicted to have a 51-amino acid deletion in the extracellular domain (PD-L1^Δ51^) overlapping Ig-like C2 domain of PD-L1.

**Figure 7.**
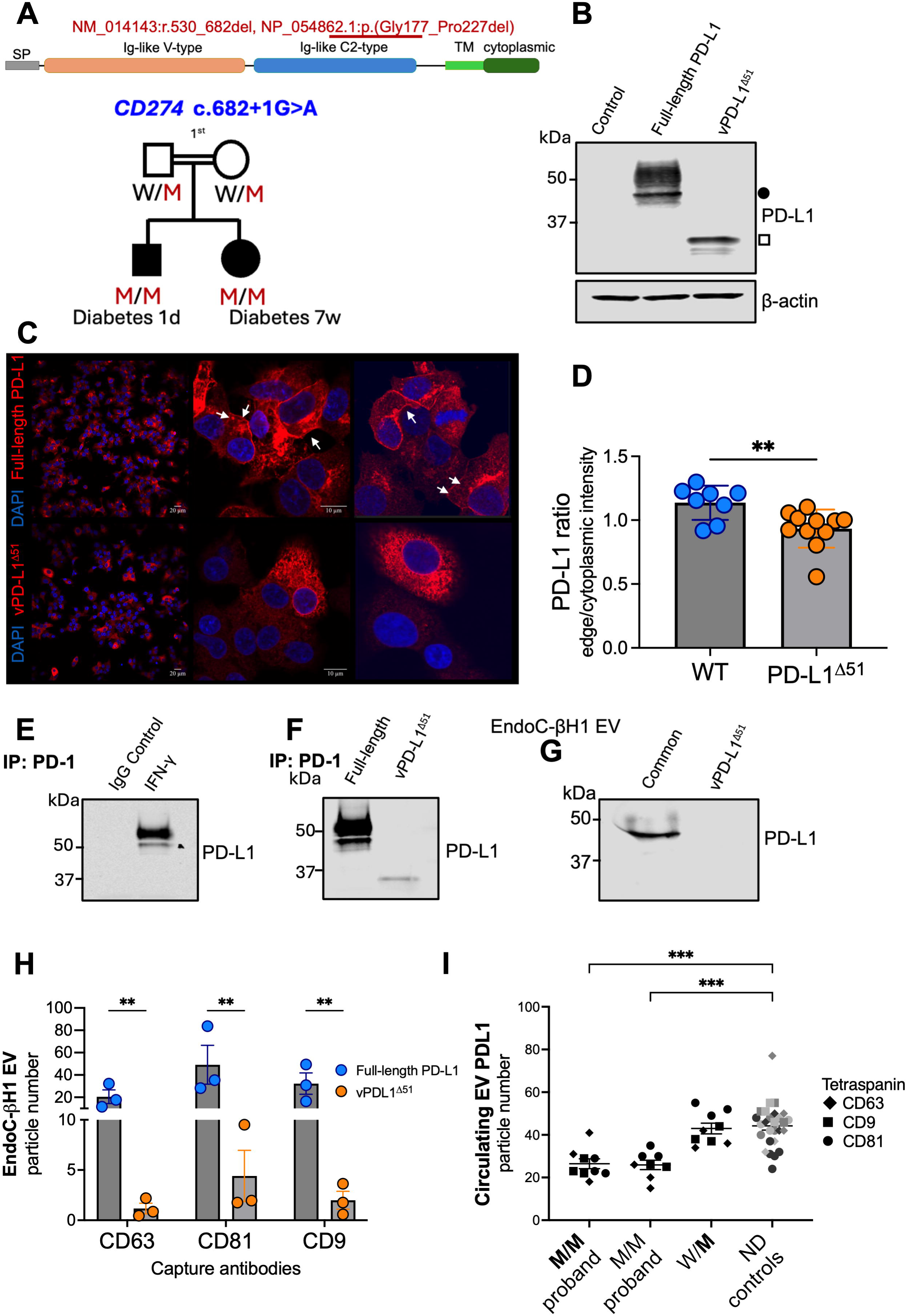
A human germline CD274 splice-site variant reproduces PD-L1^Δ3^ features and reduces circulating EV PD-L1. (A) Pedigree of a family carrying a CD274c.682+1G>A splice-donor variant. Filled symbols denote individuals with neonatal autoimmune diabetes. P1 and P2, homozygous probands (M/M); mother, heterozygous (W/M) and a schematic of resulting PD-L1 variant, red line denotes the deletion. (B) Representative immunoblot for PD-L1 and β-actin in HEK293 cells transfected with plasmid encoding full-length or variant PD-L1 (vPD-L1^Δ51^). Black circle, full-length PD-L1; white square, variant PD-L1. (C) Representative immunofluorescence images of EndoC-βH1 cells expressing full-length or variant PD-L1, stained for PD-L1 (red) and DAPI (blue). (D) Per-cell quantification of immunofluorescence images of EndoC-βH1 cells (N=3 experiments). (E) PD-1 immunoprecipitation from EndoC-βH1 cells treated with IFN-γ (50 ng/ml), followed by immunoblot for PD-L1; IgG served as control. (F) PD-1 immunoprecipitation from EndoC-βH1 cells expressing full-length or variant PD-L1. (G) Immunoblot for PD-L1 in EVs isolated from EndoC-βH1 cells expressing full-length or variant PD-L1. (**H**) Single-particle fluorescence interferometry (ExoView) quantification of PD-L1-positive EVs captured on CD63, CD81, or CD9. 3 independent experiments. (I) Circulating EV PD-L1 particle counts in plasma from homozygous probands (M/M P1, M/M P2), the heterozygous mother (W/M), non-diabetic control children (ND, N = 3, each donor in shade of gray), captured on CD63, CD9, or CD81. * P <0.05, ** P<0.01, *** P <0.001.

To characterize impacts of the resulting variant protein, we expressed the full-length (reference) or variant PD-L1 sequence in human cells. In HEK293 cells transfected with plasmid encoding the full-length or variant PD-L1^Δ51^, variant protein produced a lower-molecular-weight PD-L1 species (∼37 kDa) distinct from the full-length isoform (∼50 kDa), confirming generation of a stable, truncated protein (Figure 7B). In human EndoC-βH1 cells transfected with mRNA-loaded lipid nanoparticles (LNP) as previously described^36^, the full-length PD-L1 isoform localized to the plasma membrane, whereas the PD-L1^Δ51^ was retained intracellularly with diffuse cytoplasmic staining (Figure 7C-D), mirroring the mislocalization of stress-induced PD-L1^Δ3^ in Figure 4. Consistent with reduced PD-1 binding, PD-1 immunoprecipitation recovered IFN-γ-induced endogenous PD-L1 and LNP-delivered full-length PD-L1, but not PD-L1^Δ51^ (Figure 7E-F).

We next asked whether the variant, like PD-L1^Δ3^, is excluded from EVs. EVs isolated from EndoC-βH1 cells expressing full-length PD-L1 carried abundant full-length PD-L1, whereas EVs from variant-expressing cells were markedly depleted of PD-L1 by immunoblot (Figure 7G). ExoView across EV subpopulations confirmed a significant reduction in the percentage of PD-L1-positive particles from cells overexpressing PD-L1^Δ51^ relative to full-length PD-L1 (Figure 7H). Finally, to establish *in vivo* relevance, we quantified circulating EV PD-L1 in plasma samples from two siblings homozygous for the variant PD-L1^Δ51^ (M/M) probands who developed neonatal autoimmune diabetes^35^, as well as their clinically unaffected heterozygous (W/M) mother, and a small group of non-diabetic (ND) children and children with early onset T1D as comparators. Both of the homozygous siblings had significantly fewer EV PD-L1 particles compared to non-diabetic controls across all three tetraspanin captures. The heterozygous parent exhibited values similar to controls. (Figure 7I). Together, these human data demonstrate that a germline *CD274* splice-site variant reproduces the defining features of stress-induced PD-L1^Δ3^, resulting in intracellular retention, loss of PD-1 binding, and EV exclusion, and reduced circulating EV PD-L1 in affected individuals. These findings provide human genetic evidence that disrupted PD-L1 splicing compromises the EV-competent, immunoregulatory PD-L1 pool.

## Discussion

Here we identify PD-L1^Δ3^, an alternatively spliced isoform of PD-L1 that lacks cassette exon 3 and is generated in human β cells and islets in response to IFNs and CVB infection. PD-L1^Δ3^ is retained intracellularly rather than displayed at the plasma membrane and demonstrates diminished binding capacity to its receptor PD-1. Consistent with these properties, PD-L1^Δ3^ fails both to suppress T cell responses and to load into EVs. Elevated islet PD-L1^Δ3^ is associated with AAB^+^ and T1D, confirming disease relevance. Finally, individuals harboring a mutation yielding alternative splicing in PD-L1 exhibit similar findings to our in vitro models, resulting in neonatal autoimmune diabetes and reduced circulating EV PD-L1. These findings reframe IFN-induced PD-L1 regulation, suggesting that the β cell response to IFNs also includes a splicing shift that increases production of an isoform that is neither trafficking-nor binding-competent. Relative shifts in PD-L1 splicing could ultimately impact the balance of immune activation and β cell survival in the context of islet IFN signaling.

Prior work has identified increases in alternative splicing of other transcripts in the context of T1D development^15,16,21,23^. In the case of PD-L1^Δ3^, the exonic architecture is central to interpreting its function^27–29^. Because exon 3 encodes part of the immunoglobulin-variable domain that mediates PD-1 engagement, its loss abolishes PD-1 binding, distinguishing PD-L1^Δ3^ from full-length PD-L1 not only in localization but in intrinsic receptor affinity. This dual deficit provides a mechanistic explanation for why increased total PD-L1 transcript or protein in stressed or diabetic islets may not translate into effective immune restraint; a meaningful fraction of that PD-L1 may be both sequestered intracellularly and incapable of effective signaling through PD-1. It also underscores the importance of distinguishing isoforms, rather than measuring total PD-L1, when assessing checkpoint capacity in the β cell. Because commonly used antibodies and primer sets detect epitopes and sequences shared by both isoforms, previous reports of increased PD-L1 in T1D islets are likely to have measured the sum of full-length PD-L1 and PD-L1^Δ3^. Total PD-L1 may therefore be a poor surrogate for functional checkpoint capacity, and isoform-resolved measurement or measurement of the EV-competent pool, which is composed almost exclusively of full-length protein may better reflect the β cell’s actual capacity to restrain T cells. PD-L1^Δ3^ could also theoretically function as a dominant-negative by sequestering full-length PD-L1 and competing for shared trafficking machinery. However, a classical dominant-negative effect would demonstrate reduced CD8^+^ and CD4^+^-T cell suppression below the endogenous baseline, which we did not observe. Such interference could potentially affect EV sorting, where Δ3 might impair loading or export of full-length PD-L1 without abolishing the modest contact-dependent activity retained by the overexpressing cell. Distinguishing these possibilities will require further testing.

The two IFNs differed in their effects on the isoform balance: IFN-□ induced full-length *CD274* substantially more than IFN-α, whereas induction of *CD274*^Δ*3*^ was higher in IFN-α compared to IFN-□ (Figure 1D-E). This indicates that exon 3 skipping is not simply proportional to *CD274* transcription and raises the possibility that the relative contribution of type I versus type II IFN signaling shapes isoform balance within the islet. Based on this, we performed a time course experiment in which PD-L1^Δ3^ transcripts accumulated more rapidly than full-length PD-L1 following IFN-α exposure. One interpretation is that an early bias toward the non-suppressive PD-L1^Δ3^ isoform represents a feature of the normal antiviral response, favoring immune recognition and clearance of infected β cells, with full-length PD-L1 upregulated later in the response to restore immune restraint once the infectious threat is contained. Under these circumstances, a transient early bias toward the non-suppressive PD-L1^Δ3^ isoform would be adaptive. However, sustained increases in PD-L1^Δ3^ or relative shifts in isoform balance, such as those occurring in the described siblings with impaired splicing, could lead to unchecked β cell autoimmunity. This relationship is consistent with our observed association between CD3^+^ infiltrating T cells and islets with high β cell PD-L1^Δ3^ content in a donor with T1D and insulitis. Notably, 1 of 3 single autoantibody-positive donors showed a high relative PDL1^Δ3^ fraction, suggesting that isoform balance can shift in the absence of clinical diabetes. Because β cell IFN signaling is pharmacologically targetable, isoform-resolved PD-L1 measurement could serve as a pharmacodynamic readout of target engagement for such agents.

Our EV findings position isoform-selective cargo sorting as a decisive step in β cell–immune communication. Full-length PD-L1 is efficiently loaded onto β cell EVs, whereas PD-L1^Δ3^ is largely excluded, indicating that membrane sorting machinery discriminates between isoforms that differ only by a single internal exon. Future work will define molecular regulators of this sorting. These results also carry practical translational implications. A requirement for membrane trafficking for subsequent sorting into EVs suggests that EV PD-L1 reflects full-length PD-L1 at the cell membrane and could serve as a readout of IFN signaling. The observations that subsets of Ab+ or T1D donors showed relative increases in PD-L1^Δ3^ compared to full-length PD-L1, and that PD-L1^Δ3^ was observed in β-cells near CD3+ infiltrates, further suggests that isoform balance could underlie inter-individual heterogeneity in β cell immune signaling. Because only full-length PD-L1 is effectively packaged into EVs, EV PD-L1 could serve as a noninvasive readout of relative isoform balance.

A germline *CD274* splice-site variant provides convergent, human genetic support for this model. Although the c.682+1G>A variant and IFN-induced exon 3 skipping are mechanistically distinct events, they present similar phenotypes, indicating that the integrity of CD274 splicing, not merely its expression level, governs the potential for PD-L1’s immunoregulatory function and export into EVs. Notably, this phenotype implicates the β cell as a highly vulnerable component of this axis. Despite PD-L1’s broad expression across immune lineages, the homozygous siblings showed largely normal lymphoid and myeloid development, unlike the extensive immune dysregulation found in inherited PD-1 deficiency, yet both developed neonatal onset T1D^35^. This sparing of the immune compartment alongside early, penetrant β cell autoimmune destruction argues that disease is driven less by loss of PD-L1 on immune cells than by loss of membrane- and EV-competent PD-L1 on the β cell itself, depriving it of a cell-intrinsic means of restraining infiltrating T cells. Stress-induced PD-L1^Δ3^ may thus phenocopy, in an acquired and potentially reversible fashion, the β cell-intrinsic vulnerability imposed constitutively by the germline variant.

Several limitations warrant mention. The modest CD8^+^ and CD4^+^-T cell suppression by PD-L1^Δ3^ overexpressing cells suggests residual activity, either from a small surface-competent fraction of overexpressed protein or from PD-1-independent immunomodulatory effects of the retained pool. The trans-acting factors that couple inflammatory signaling to exon 3 skipping and to isoform-selective EV loading remain to be defined. Addressing these questions will clarify whether the PD-L1^Δ3^ splicing shift is a maladaptive failure of an otherwise protective program or a regulated tuning of β cell immune signaling and whether it can be therapeutically redirected toward EV-competent, immunoregulatory PD-L1. Pancreas sections from donors with AAB+ or T1D are cross-sectional, and so we are unable to examine relationships with diabetes progression or residual insulin secretion. The number of donors with AAB^+^ and T1D are small, limiting the power of spatial comparisons. Longitudinal testing in natural history cohorts will be required to understand relationships of relative increases in circulating EV PD-L1 with disease natural history.

In summary, we define a post-transcriptional axis in which β cell interferon signaling not only increases transcription of full-length PD-L1, but also diverts β cell PD-L1 transcription toward an intracellularly retained, PD-1 binding-deficient, EV-excluded isoform with reduced functional capacity to impact T cells. This work establishes isoform balance and EV partitioning of PD-L1 as previously unrecognized determinants of β cell–immune communication in T1D and nominates them as candidate biomarkers of β cell interferon signaling and potential contributors to heterogeneity in the natural history of β cell survival.

## Research Design and Methods

### Sex as a biological variable

Human islet, pancreatic tissue, and blood samples were obtained from both male and female donors, based on availability, and analyses combined data from both sexes. Findings are expected to be relevant to both sexes; the study was not powered to detect sex-specific differences.

### Human islets

Deidentified nondiabetic male and female human donor islets were obtained from the Integrated Islet Distribution Program (IIDP) or the University of Alberta Diabetes Institute Islet Core (Edmonton, Alberta, Canada) or the United Network for Organ Sharing (UNOS) (see Table 1). Deidentified paraffin-embedded pancreatic tissue sections from donors with T1D, single autoantibody positive (sAAb⁺) donors, and non-diabetic controls were obtained from the Network for Pancreatic Organ Donors with Diabetes (nPOD) (see Table 2). The use of deidentified human samples was approved by the Institutional Review Board at the Indiana University School of Medicine and considered exempt from human subjects research.

### Cell culture

βlox5 cells, a gift from Dr. Jon Piganelli, were cultured in RPMI medium (Gibco) containing 10% heat-inactivated FBS (Gibco; S11550H, Lot-A20005) and supplemented to a final concentration with L-glutamine (Thermo Scientific, 2 mM), penicillin (Corning, 50 U/ml), streptomycin (Corning, 50 μg/ml), HEPES (1M Thermo Scientific; 15630080), Sodium pyruvate (1%; Thermo Scientific), NEAA (1%; Thermo Scientific; 11140-050). EndoC-βH1 cells, obtained from Human Cell Design (Toulouse, France) were cultured in low glucose DMEM (5.5mM) supplemented with 2% BSA, 50μM β-mercaptoethanol, 10 mM nicotinamide, 5.5 μg/ml transferrin, 6.7 ng/mL sodium selenite, and 1% P/S, in plates pre-coated with matrigel-fibronectin. HEK293 cells were cultured in high glucose DMEM supplemented with 10% FBS, 1% P/S, and 1% L-Glutamine. Human islets were cultured in standard islet medium (Prodo; PIM-S001GMP), supplemented with human AB serum (Prodo; PIM-ABS001GMP), Glutamine and glutathione (Prodo; PIM-G001GMP), and ciprofloxacin (Fisher; MT61277RG).

### Cytokine treatment

Cells and human islets were treated with 2000 U/ml IFN-α2a (PBL Assay Science; 11100) or 100 ng/ml human IFN-γ (R&D systems; 285IF100) for 24 h unless otherwise indicated.

### Enterovirus infection

Human islets (400 islet equivalents per condition) were treated with vehicle control or infected with human Coxsackievirus B3 (CVB3, MOI = 50; ATCC® VR-30™). At 24 h post-infection, islets were UV crosslinked to inactivate virus and subsequently fixed in 4% paraformaldehyde on ice for 20–30 min. Fixed islets were washed with PBS and cryoprotected overnight at 4°C in 30% sucrose (w/v) in PBS. Islets were then washed, embedded in optimal cutting temperature (OCT) compound, and frozen on dry ice.

### Reverse transcriptase PCR and qPCR

RNA stored in Qiazol (Qiagen; 79306) was extracted using RNeasy Plus Mini kit (Qiagen; 74134), and cDNA synthesis was performed using iScript cDNA Synthesis Kit (BioRad; 1708890) according to manufacturer’s instructions. RT-PCR was performed with primers flanking exon 2-exon 4 to resolve full-length (500 bp) and exon 3-skipped (184 bp) products (full-length, F: ATGGTGGTGCCGACTACAAG, R: GGAATTGGTGGTGGTGGTCT; exon-3 skipped, F: TTGCTGAACGCCCCATACAA, R: TCCAGATGACTTCGGCCTTG); products were resolved on a 1% agarose gel. Isoform-specific qPCR was performed using TaqMan assays (Applied Biosystems; A58666) targeting full-length CD274 (assay ID: Hs.PT.58.41014847) and the exon 2-exon 4 junction of CD274^Δ3^ (assay ID: Hs.PT.58.46567069). A relative standard curve was generated on each plate from serial dilutions of a pooled cDNA reference (0.2-100 ng per reaction), and transcript levels were interpolated from this curve (R² > 0.99, amplification efficiency 90%-110%). Values were normalized to ACTB (assay ID: Hs.PT.39a.22214847) and are reported as relative quantity (RQ) in arbitrary units; time-course data are expressed as fold change from 0 hours.

### PD-L1 isoform overexpression

Human full-length PD-L1 (EX-U0767-LX304-B), FLAG-tagged PD-L1^Δ3^ (EX-A1981-Lv242-B), empty control vector (EX-NEG-Lv242-B) were purchased from Genecopoeia. Constructs were packaged into lentiviral particles using HEK 293T cells co-transfected with viral packaging plasmids, psPAX2 and pMD2.G (Addgene; 12260, 12259) as previously described^37^. Lentiviral supernatants were harvested after 72 h, and βlox-5 cells were transduced with filtered lentivirus and selected using puromycin (5 μg/ml, InvivoGen; ant-pr-1).

For select experiments, mRNA encoding PD-L1 or PD-L1^Δ51^ was delivered using a lipid nanoparticle (LNP) platform as previously described^36^. mRNA for LNP delivery was synthesized by Genscript based on published sequences for PD-L1 and PD-L1^Δ51^ variant^35^. EndoC-βH1 cells were transfected with LNP-encapsulated mRNA at 1.5 µg mRNA per well in a 6-well plate for 18 hours prior to treatments as described in each figure. Transduction efficiency was confirmed using LNPs encapsulating mScarlet mRNA. For experiments in HEK293 cells, plasmids encoding C-terminally FLAG-tagged full-length PD-L1 or PD-L1^Δ51^ were transfected using Lipofectamine 3000 (Thermo Fisher Scientific) according to the manufacturer’s instructions.

Presence of PD-L1, PD-L1^Δ3^, or PD-L1^Δ51^ overexpression in the cells was confirmed by immunoblot.

### BaseScope Assay

Isoform-specific BaseScope probes were designed to distinguish full-length PD-L1 (targeting exon 3; ACD 1118181-C1) from PD-L1^Δ3^ (targeting the exon 2-exon 4 junction; ACD 1242441-C1). For paraffin embedded tissue, the slides were baked at 60°C for 1 hour using hybEZ oven (ACD; 321710/321720) and then the tissue sections were deparaffinized by incubating the slides twice with xylene for 5 min and 100% ethanol for 2 min at RT. The slides were air dried at 60°C for 5 min, then moved to RT, and treated with hydrogen peroxide (ACD) for 10 min. After washing 2X with ddH2O, the slides were moved to a container with 1X antigen-retrieval buffer (ACD) at 99-100°C for 20 min. Next, slides were washed with ddH2O for 15 secs at RT, followed by incubation with 100% ethanol for 3 min and air-drying at RT. Sections were applied with a hydrophobic barrier (Immedge, Vector labs) and air-dried at RT. The slides were transferred to a humidifying chamber and treated with protease III (ACD) for 20 min at 40°C in a HybEZ Oven (ACD). Next, slides were washed 5X with ddH2O, and hybridization was performed for 2 hours using BaseScopev2 assay probe (ACD, 1118181-C1, 1242441-C1). Two control probe sets, PPIB-1zzas (18282A, positive control) and bacterial DapB-1zz (18267A, negative control), were used as internal controls to determine assay specificity. Signal was developed using BaseScope Reagent kit v2 RED (ACD; 323900) according to the manufacturer’s instructions.

Frozen 8-μm sections containing human islets CVB3 were fixed in 10% neutral buffered formalin for 15 min at 4°C, and dehydrated through 50%, 70%, and 100% ethanol (5 min each at RT). Sections were then pretreated with hydrogen peroxide and hybridized as described above.

After hybridization, slides were washed twice in ddH₂O and twice in PBST, blocked in donkey serum in PBS for 10 min, and immunostained for human insulin (Dako; A0564) or human CD3 (Dako; A0452) as previously described. Nuclei were stained with DAPI (ACD), and slides were mounted in ProLong Gold Antifade medium (Invitrogen). Imaging was performed on a Zeiss LSM800 confocal microscope or Keyence BZ-X1000 using a 40x oil objective or 10x objective with Z-stacks.

### Image quantification (QuPath)

BaseScope signal was quantified in QuPath v0.6.0. Images were opened via Bio-Formats and set to fluorescence type, with channels assigned as insulin (Alexa 488), BaseScope (Texas Red), and DAPI. Islets were defined by insulin positivity: a pixel classifier/thresholder was applied to the insulin channel, thresholded regions were converted to annotations, and annotations were manually reviewed to merge or split over- and under-segmented islets, yielding one annotation per islet. Islets at tissue edges, those with >50% of their area outside the field, and those with visible artifactual autofluorescence were excluded. BaseScope puncta weren detected within islet annotations using subcellular spot detection on the Texas Red channel with an expected spot size of 0.3-0.6 μm and a fixed intensity threshold applied uniformly across all slides in a batch; detection was restricted to islet annotations. Per-islet spot counts and islet area (μm²) were exported as CSV, and transcript density was calculated as spots per 1000 μm² of insulin-positive islet area. Quantification was performed blinded to donor group.

### Protein isolation and immunoblotting

Protein isolation and immunoblotting were performed as previously described^14^. Briefly, whole cell extracts of cells were prepared in a lysis and extraction buffer (Thermo Scientific; 89901) supplemented with HALT protease inhibitor cocktail (Thermo Scientific; 78430). Cytoplasmic, membrane, and nuclear fractions were prepared using the Subcellular Protein Fractionation Kit for Cultured Cells (Thermo Scientific; PI78840) according to the manufacturer’s instructions, and fraction purity was confirmed by immunoblot. Samples were diluted using 1X sample buffer (LI-COR Biosciences; 928-40004) with 2-Mercaptoethanol (Thermo Scientific; 21985023). The proteins were separated using 4-20% SDS-PAGE precast gel (Bio-Rad; 4561094) and transferred to activated PVDF membrane and membranes were blocked with Intercept® (TBS) blocking buffer (Li-Cor Biosciences) for 1 h. The blots were probed with the following primary antibodies and 0.2% Tween20 with overnight incubation at 4°C; anti-PD-L1 (Cell signaling;13684; 1:750) (Cell Signaling; 29122; 1:1000), anti-CD63 (Antibodies Online; ABIN144001; 1:1000), anti-DYKDDDDK Tag (Cell Signaling; 14793S; 1:500), anti-tubulin (Cell Signaling; 2125S; 1:2000), anti-E-cadherin (Cell signaling; 14472; 1:1000), anti-Lamin A/C (Cell signaling; 4777; 1:1000), anti-CD9 (Proteintech; 20597-1-AP; 1:1000), anti-CD81 (Invitrogen; SN206-01; 1:1000), anti-Calreticulin (Proteintech; 27298-1-AP; 1:1000) and anti-β actin (Cell signaling; 4970S; 1:1000). Anti-rabbit or anti-mouse IgG (H+L)-HRP conjugate, IRDye 680-conjugated, or IRDye 800-conjugated (Li-Cor BioSciences; 1:10,000) secondary antibodies were used for visualization and quantification. Immunoblots were visualized using the Li-Cor Odyssey system (Li-Cor Biosciences) and quantitated using Odyssey Imaging software (Li-Cor Biosciences).

### Flow cytometry

For surface PD-L1 staining, βlox5 cells expressing empty vector, full-length PD-L1, or PD-L1^Δ3^ were dissociated with 0.05% trypsin, washed in FACS buffer (PBS with 2% FBS), and stained with anti-PD-L1 (clone 28-8; Abcam; ab206967) or isotype control for 30 min at 4°C. Cells were washed, stained with a fixable viability dye, and acquired on an Attune NxT flow cytometer (Thermo Scientific). Data were analyzed in FlowJo v10 (BD Biosciences), and PD-L1 mean fluorescence intensity was determined on live single cells.

### PD-1/PD-L1 interaction assay

Recombinant His-tagged human PD-1 (R&D Systems; 8986-PD) was immobilized on Ni-NTA magnetic beads (Thermo Scientific; 88831) according to the manufacturer’s instructions. PD-1 bound beads were added to cultured PD-L1-expressing cells and incubated for 4 hours at 37°C. Cells were lysed in standard lysis buffer, and bead-bound complexes were washed and eluted in 25 mM Tris-HCl (pH 7.5), 10 mM NaCl, 0.1% SDS, and 100 mM dithiothreitol. Eluates were immunoblotted for PD-L1 and FLAG. Beads without immobilized PD-1 served as negative controls.

### Human T-cell coculture assay

Peripheral blood mononuclear cells were purchased from ImmunuSpot (CTL, Table 3) and cultured in DMEM media with 10% FBS, L-Glut, NEAA, HEPES, Sodium pyruvate and 2-Mercaptoethanol. Following 24-48 h, the cells were labeled with CellTrace Violet (CTV, Invitrogen; C10094) according to manufacturer’s protocol and stimulated with Concanavalin-A (Con-A, Sigma; C5275). CTV labelled PBMCs were co-cultured with β cells expressing full-length PD-L1 or PD-L1^Δ3^ at a 1:1 effector-to-target ratio and analyzed by flow cytometry at 24 h, 48 h and 72 h time points. Cells were also stained with eFluor 780 fixable viability dye (eBiosciences; 650865) and to assess CD8+ and CD4+ T cell proliferation, expression of classic early activation marker (CD69) and inhibitory/checkpoint receptor (PD1), using the following primary antibodies CD4 (Biolegend; 344606), CD8 (Biolegend; 344726), CD69 (Biolegend; 310938), PD-1 (Biolegend; 367428). Culture supernatant fraction was collected and stored at −80°C for cytokine analysis. Samples were recorded on an Attune NxT Flow Cytometer (Thermo Scientific) and data were analyzed using FlowJo v10 Software (BD Biosciences). PBMC coculture supernatant fractions were assayed by IFN-γ ELISA (BioLegend; 430104). ELISAs were performed according to manufacturers’ protocols and read on a SpectraMax M2 microplate reader (Molecular Devices). All samples were tested in duplicate.

**Table 3.** Human PBMC characteristics.

| **Donor No** | **SampleID** | **Age** | **Sex** | **Source** |
| --- | --- | --- | --- | --- |
| 403 | HHU20200218 | 28 | Male | CTL ePBMC |
| 406 | HHU20200305 | 54 | Male | CTL ePBMC |

**Table 4.**
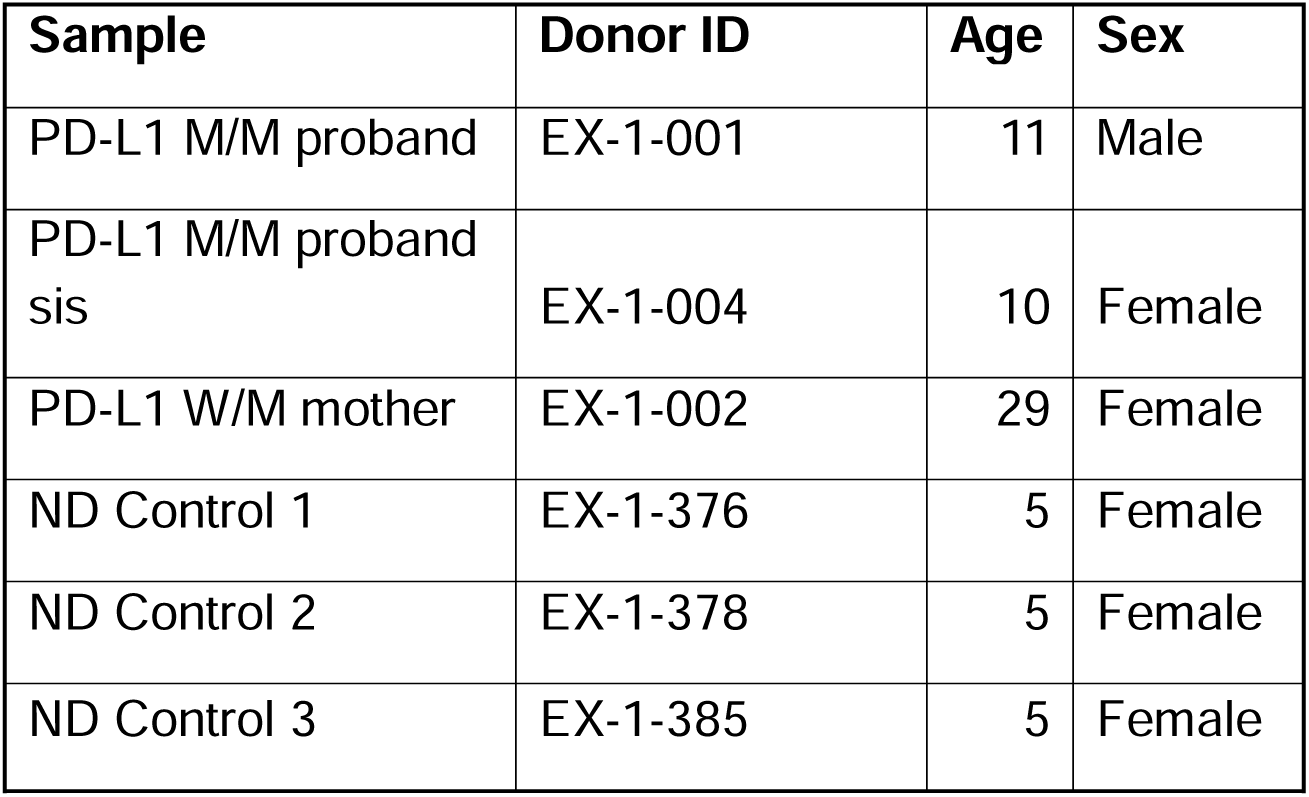
Human Plasma characteristics.

| **Sample** | **Donor ID** | **Age** | **Sex** |
| --- | --- | --- | --- |
| PD-L1 M/M proband | EX-1-001 | 11 | Male |
| PD-L1 M/M proband<br>sis | EX-1-004 | 10 | Female |
| PD-L1 W/M mother | EX-1-002 | 29 | Female |
| ND Control 1 | EX-1-376 | 5 | Female |
| ND Control 2 | EX-1-378 | 5 | Female |
| ND Control 3 | EX-1-385 | 5 | Female |

### EV isolation and characterization

For cell lines, 10-15 ml of supernatant was centrifuged at 800 g for 10 min. The supernatant fraction was centrifuged at 2,000 g, passed through a 0.22 µm filter (MerckMillipore; GSWP02500) and was centrifuged at 100,000 g for 1.5 h at 4°C to pellet small EVs. Total EV protein concentrations were determined by using a BCA protein assay kit (Thermo Scientific; 23225). EVs were isolated from human islet medium and 150 µl plasma using size exclusion chromatography (SEC). Samples were centrifuged at 2,000 g for 10 min and ultrafiltered using 0.45 µm filter (Cytivia; 6780-2510). After rinsing the qEV single columns (IZON Science; ICS-70) with 0.22 µm filtered 1X PBS, 500 µl of the sample was applied on top of a column and 0.5 ml fractions were collected in 1.5 ml tubes. Four EV-rich fractions (6-9) were pooled and analyzed for EV purity.

### Single-particle interferometry (ExoView)

EVs were captured on chips bearing anti-CD63, anti-CD81, and anti-CD9 antibodies and probed for PD-L1 (clone 28-8; Abcam; ab206967) (R&D Systems; FAB1562R) using the ExoView platform (Unchained Labs; 251-1064) according to the manufacturer’s instructions. Particle counts were analyzed using ExoView Analyzer software.

### Dot Blot

Eluted SEC fractions were blotted (2 μL per fraction) onto 0.2 μm nitrocellulose membranes (Thermo Scientific, 77012) and allowed to dry for 30 min. Membranes were blocked with Intercept Blocking Buffer (Li-Cor Biosciences, 927-60001) for 1 hr at RT and then incubated with primary antibodies overnight at 4°C. Membranes were washed four times for 5 min with TBS supplemented with 0.2% TBST at RT before incubation with secondary antibodies for 45 min at RT. Membranes were washed four times for 5 min with TBST prior to detecting fluorescence signals using the Li-Cor Odyssey CLx imaging system (Li-Cor Biosciences).

### Variant PD-L1^Δ51^ immunofluorescence

EndoC-βH1 cells transduced with mRNA LNP containing PD-L1 or PD-L1^Δ51^ were fixed in 4% paraformaldehyde for 15 min, permeabilized in 0.5% saponin for 15 min, and blocked in 1% BSA in PBS containing 0.1% Tween-20. Samples were incubated with anti-PD-L1 (Cell Signaling; 29122S, 1:100 dilution) for 1 h at room temperature, followed by AlexaFluor-conjugated secondary antibody labeling for 1 h (Invitrogen), and counterstained with DAPI. Images were acquired on a Nikon A1 confocal microscope (Nikon) and analyzed using CellProfiler.

### Human Plasma Samples (Exeter)

EDTA blood samples from family members carrying the CD274 c.682+1G>A variant and from control participants were collected at the University of Exeter, UK after written informed consent. Plasma was separated from the EDTA whole blood following centrifugation at 12,000g. The study was approved by the East Midlands – Derby Research Ethics Committee (17/EM/0255, IRAS 228082).

### Quantification and statistical analysis

All data are represented as mean ± SEM. For comparisons involving more than 2 conditions, 1-way ANOVA or repeated-measures ANOVA (with Tukey’s post hoc test or Dunnett’s post hoc test) was performed. For comparisons involving only 2 conditions, a 2-tailed Student’s unpaired *t* test was performed. GraphPad Prism, version 11, was used for all statistical analyses and visualization. Statistical significance was assumed at *P* < 0.05.

## Acknowledgements

Human pancreatic islets were provided either by the NIDDK-funded Integrated Islet Distribution Program (IIDP) (RRID:SCR _014387) at City of Hope, NIH Grant # U24DK098085 and the JDRF-funded IIDP Islet Award Initiative, or provided by the Alberta Diabetes Institute IsletCore at the University of Alberta in Edmonton (http://www.bcell.org/adi-isletcore.html) with the assistance of the Human Organ Procurement and Exchange (HOPE) program, Trillium Gift of Life Network (TGLN), and other Canadian organ procurement organizations. Islet isolation was approved by the Human Research Ethics Board at the University of Alberta (Pro00013094). All donors’ families gave informed consent for the use of pancreatic tissue in research.

This research was performed with the support of the Network for Pancreatic Organ donors with Diabetes (nPOD; RRID:SCR_014641), a collaborative type 1 diabetes research project supported by Breakthrough T1D and The Leona M. & Harry B. Helmsley Charitable Trust (Grant#3-SRA-2023-1417-S-B). The content and views expressed are the responsibility of the authors and do not necessarily reflect the official view of nPOD. Organ Procurement Organizations (OPO) partnering with nPOD to provide research resources are listed at https://npod.org/for-partners/npod-partners/.

## Author contributions

CR, EKS conceptualized the research. CR, FH, SR, MCA, JDP, AKL, RGM, EKS designed research studies. CR, FH, SR, MCA, IA, AGO acquired data. CR, FH, SR, KK analyzed the data. JDP, MBJ and RO provided the reagents. MBJ, DLE, CEM, AKL, JDP, RGM and EKS provided project supervision. CR and EKS wrote the original draft. All authors contributed to discussion, edited the manuscript, and approved the final version of the manuscript.

